# Development of a new tool based on gene expression analyses of bovine raw milk to monitor the inflammatory status of the udder

**DOI:** 10.1101/2025.09.03.671445

**Authors:** C. Gitton, Y. Le Vern, M. Gaborit, R. Prado Martins, P. Germon

**Author notes:** PFIE UE 1277, INRAE, Nouzilly, France.

## Abstract

Inflammation of the mammary gland, or mastitis, is a common disease of dairy cows with important consequences on the economy of dairy farms, well-being of cows and antibiotic usage. Monitoring of the inflammatory status of mammary gland currently relies on milk somatic cell counting. In order to improve this monitoring, methods aiming at differentiating cell types among milk cells are being developed. The objective of the present study was to explore the use of gene expression profiling from raw milk samples to define the inflammatory status of udder quarters. A pilot study allowed us to select 37 genes differentially expressed between three milk cell types, i.e. neutrophils, macrophages or T lymphocytes. Based on expression data measured by RT-qPCR for these genes in RNA extracted from quarter milk samples, we were able to identify four different types of milk samples, each with specific properties. One cluster was characterized by increased expression of genes typifying neutrophils, a second one was identified by increased expression of genes typifying macrophages while the two others were distinguished based on the expression of genes associated with T lymphocytes. Clustering of samples based on gene expression was shown to be comparable to that obtained by flow cytometry analysis. Our results highlight the potential of profiling of milk samples based on gene expression for the analysis of the inflammatory status of udders.

## Introduction

Mastitis remains a major issue in dairy farming. Upon entry and recognition of bacterial pathogens by the host immune cells, in particular mammary epithelial cells and macrophages, an inflammatory response is triggered which leads to recruitment of immune cells in the mammary gland.

As a consequence, somatic cell counts (SCC) in milk is routinely used as an indirect indicator of the sanitary status of mammary glands.

SCC values below 100,000 cells per ml are generally considered as an indication of a healthy mammary quarter (1, 2). On the opposite, SCC values above 200,000 cells/ml are generally taken as an indication of an on-going inflammatory response or a quarter recovering from infection (2, 3). In between these two SCC values one cannot state with confidence whether a quarter is healthy or not. In addition, setting SCC threshold values to distinguish healthy from infected quarters should also take into account different criteria such as the lactation stage and parity (4). In any case, these two values are not definite indicators of a health status: mastitic quarters below SCC values of 100,000 could be still be infected by mastitis pathogens (5).

As a consequence, several improvements have been tested, in particular the differential cell counts that determines the type of cells present in milk upon inflammation (6, 7). Cells present in milk belong to different cell types including neutrophils, macrophages, lymphocytes and epithelial cells. The proportion and quantities of each of these cell population is highly variable. For instance, the strong inflammation observed during clinical mastitis is characterized by a majority of neutrophils which can amount to almost 90% of cells present in milk (8).

Reports on the cell composition of mammary gland secretions from healthy animals show contrasting results. While several studies revealed mostly macrophages (50-80%) with 10-20 % lymphocytes (9-13), others rather indicate that T lymphocytes are the most prevalent cell type in the absence of infections with almost 50% T lymphocytes and around 20-30 % macrophages (14-16).

These discrepancies might be due to different methodologies, to the potential confusion between macrophages and epithelial cells after staining and light microscopy examination (17). The cell composition is also highly dependent on the inflammatory status of the mammary gland, with increased prevalence of neutrophils and decreasing percentages for lymphocytes, and on the lactation status with increased values in late lactation for macrophages (15, 17).

Flow cytometry analyses identified different subpopulations of T lymphocytes, in particular CD4+, CD8+ as well as *γδ* T cells, along with different macrophage classical (CD14+ CD16-) and non-classical (CD14-CD16+) subpopulations (18). While CD4 and CD8 T lymphocytes constitute approximately 20-30 % of all T-cells, *γδ* T cells are also present in milk in significant numbers (10-30% of all lymphocytes in milk) (17-19). The proportion of these subpopulations among T lymphocytes depends on the mammary gland health status: indeed, the CD4/CD8 ratio increases upon infection or inflammation (14, 20-23) as well as in late lactation (19). While B-cells are only present in limited numbers in healthy mammary glands, their proportion among lymphocytes was shown to increase upon inflammation (24).

Based on such evidence that the proportion of the different cell types is related to the health status of a mammary gland, differential cell count strategies have been explored in order to not only rely on SCC values to characterize diseased quarters (6, 15). One such Differential Somatic Cell Count (DSCC) method discriminates between neutrophils and macrophages while lymphocytes are counted along with neutrophils (25, 26). Four groups based on SCC and DSCC values were determined. Similar to previous studies, this method highlights variations depending on parity and lactation status (27, 28). Although both indicators are linked to the recruitment of leukocytes in mammary gland secretions, their dynamics was found to be slightly different, with DSCC values decreasing faster than SCC in the resolutive phase of inflammation (29). DSCC can thus be considered as an additional indicator for the control of mastitis. Yet, the use of DSCC will require the definition of suitable thresholds in order to improve its predictive value (27). In fact, a recent survey suggest very little added value of DSCC over SCC in terms of positive- and ngetive-predictive values for the detection of mastitis cases due to major pathogens (30). Furthermore, the definition of suitable thresholds for SCC/DSCC combinations will have to take into account a potential herd effect since authors have also observed significantly different values between different herds, showing that the herd effect could be more important that the pathogen (31).

High resolution differential cell count based on flow cytometry has also been proposed recently to better characterize the inflammation/infectious status of mammary glands (18). A more global approach has also been tested using single-cell RNAseq highlighting the presence of different cell populations and subpopulations in a healthy milk sample (32, 33). Altogether these different methods allow the identification of cell populations present in milk.

We hypothesized that another strategy to characterize the inflammatory status of mammary glands could be based on a functional assessment of the cells present in mammary gland secretion. In order to do so, we first identified a subset of genes specific for macrophages, neutrophils or T lymphocytes, three of the main cell populations present in milk. We then undertook a gene-expression analysis of these selected genes based on RNA samples collected from unprocessed raw milk. Gene expression profiles were determined allowing clustering of milk samples and the clustering obtained was then compared to results obtained by flow cytometry.

## Materials and methods

### Ethics approval

Not applicable. Milk samples were collected from cows as part of a routine milking.

### Collection of milk samples

Quarter foremilk samples (40 ml) from all four quarters were collected under aseptic conditions as recommended by the National Mastitis Council guidelines, from 28 Holstein (HF) and 10 Normande (NO) cows at the experimental farm of Le Pin-au-Haras (Unité Expérimentale du Pin-au-Haras). No criteria was applied for the selection of cows from which milk samples were collected. More specifically, after teat cleaning, the first streams of milk were discarded, premilking disinfectant solution was applied and then dried, teat apex was disinfected by scrubing with a 70% ethanol moistened gauze pad and milk was collected in 50ml tubes held at 45° angle to avoid contamination by dust materials. Milk was then transported to the laboratory at room temperature within 3h.

A 500 μL portion of each milk sample was subjected to a Fossomatic model 90 (Foss Electric, Hillerod, Denmark) to determine the milk SCC. The somatic cell score (SCS) was calculated with the formula : SCS = Log2(SCC / 100,000) + 3. One milk sample from each cow was selected, except for four cows for which two samples were collected from two different quarters: the only criteria for selection of milk samples was to have a panel of milk samples representing a large range of SCC values. These quarter milk samples originated from cows that were at different stages of lactation (days-in-milk ranging from 35 to 391 days) and with different SCC values (ranging from 17,000 to 3,603,000 cells/ml). None of the samples were from clinical mastitis cases. Details are presented in **Supplementary table S1**.

Bacteriological examination of the selected quarter milk samples was performed by plating 50 μL of milk on sheep blood agar plates (ThermoFicher Columbia agar with Sheep Blood PLUS #PB5039A) and recording bacterial colonies after 24 and 48 hours of incubation at 37°C. Basic tests (colony morphology and colour, hemolysis, Gram-staining, catalase) were performed in order to qualify the bacteriological status of milk samples. Samples were considered contaminated when more that 3 different colony types were observed.

### Isolation and labelling of milk cells

Milk cells were collected from the selected milk samples: An equal volume of 1× DPBS (reference, Sigma D8537) was added to milk samples. The diluted milk sample was left at room temperature for 15 minutes and then centrifuged at room temperature for 10 minutes at 1400g. The cream layer and the supernatant were removed and the cell pellet was then washed once in 10ml of room temperature (DPBS, 2mM EDTA, 1% horse serum) and then centrifuged 5 minutes at 500g. The cells were then resuspended in FACS buffer (DPBS, 2mM EDTA, 10% horse serum). Cells were then labelled prior to flow cytometry analysis or cell sorting.

Cells were quantified using a LUNA-FL cell counter (Logos Biosystems) according to the manufacturer’s instructions. One million cells prepared as indicated above were transferred to a 1.5ml tube, centrifuged and resuspended in FACS buffer. Cells were labeled for 30 min with primary antibodies (30 min, 4°C). Primary antibodies used in the present study were CD14-PE-AF750 (Bio-Rad, reference MCA1568P750, clone Tük4), CD45-PE (Bio-Rad, reference MCA2220PE, clone 1.11.32), G1 (Kingfisher Biotech, reference WSC0608B-100, clone CH138A) and CD3 (Kingfisher Biotech, reference WS0561B-100, clone MM1A). After washing in FACS buffer, cells were incubated (30 min, 4°C) with the secondary antibodies BV421 labelled rat anti-mouse IgM antibody (BD Biosciences, reference 562595, clone R6-60.2) and Alexa Fluor™ 488 labelled Goat anti-Mouse IgG1 (Thermo Fisher Scientific, reference A21121) in FACS buffer, matching respectively G1 and CD3 primary antibodies. After washing in FACS buffer, cells were resuspended in DPBS containing the Zombie Aqua viability dye (BioLegend, reference 423101) for 30 minutes at 4°C. After a final wash in FACS buffer, cells were resuspended in FACS buffer.

### Cells sorting of purified specific leucocytes populations

Milk cells from three quarter milk samples were collected and labelled as described above. These three samples were selected based on their SCC, representative of low, intermediate and high SCC quarters. Labelled milk cells were then sorted on a MoFlo Astrios EQ (Beckman Coulter) high speed cell sorter equipped with 4 lasers to obtain, among live cells, purified macrophages (CD45+ G1-CD14+), T lymphocytes (CD45+ CD3+) and neutrophils(CD45+ G1+). The average sorting speed was 10,000 cells/s.

### Flow cytometry analysis of milk cells

To determine the proportion of leucocyte subpopulations in milk samples, milk cells were isolated and labelled as described above. Milk cells were then analyzed on a LSR Fortessa™ X-20 Flow cytometer (Becton Dickinson) and results were analyzed with the Kaluza software v2.1 (Beckman Coulter). Among live cells, macrophages were identified as CD45+ G1-CD14+ cells, neutrophiles as CD45+ G1+ cells and T lymphocytes as CD45+ CD3+ cells. Our gating strategy is exemplified in **Supplementary Figure S1**.

### RNA extraction and high-throughput RT-qPCR

For sorted cells, immediately after collection, cells were centrifuged 10 minutes at 1000g, resuspended in 1 ml of Trizol and stored at -80°C until RNA was extracted as described below. For milk samples, 200 microliters of milk were mixed with 800 µl of Trizol (Thermofisher) in a lysing matrix D tube (MP Bio) and homogenized for 45 seconds – speed 6 in a FastPrep-24 apparatus. Samples were then stored at -80°C until further processed.

Upon thawing of milk and purified cells samples mixed with Trizol, 200 µl chloroform was added, the tube was vortexed for 10 seconds and then centrifuged at 12000g for 15min at 4°C. The aqueous phase was collected, mixed with an equal volume of 70% ethanol, loaded on a NucleoSpin RNA columns (Macherey Nagel) and RNA was then purified and treated with DNAse I as indicated by the manufacturer. Total RNA (50 ng) was reverse transcribed to cDNA using 5x iScript reverse transcription supermix (Biorad) according to manufacturer’s instructions. Ten-fold diluted cDNA were preamplified according to Fluidigm’s protocol (quick reference PN 100-5875 B1) using PreAmp master mix and a mix of all forward and reverse primers at 500nM each. Preamplification program was 2 minutes 95°C, followed by 16 cycles of (95°C for 15s, 60°C for 4 min). Preamplified cDNA were then diluted 5-fold with (10 mM Tris, pH 8.0, 1 mM EDTA) buffer. Gene expression level were measured from pre-amplified cDNA on 48.48 GE Dynamic Array IFC or 96.96 GE Dynamic Array IFC using the Fluidigm BioMark™ HD System. The full list of primers used in this study, including ones that were not retained after differential expression analysis, are listed in **Supplementary Table S2**.

### Data analysis

A reference sample was built from the arithmetic mean of Ct for each gene of analyzed samples. Fold changes compared to this reference sample were calculated by the ΔΔCt method using *ACTB* and *GAPDH* as reference genes (34). Kruskal-Wallis and Mann-Whitney tests were performed using the stats package (v4.5.1). Heatmaps were generated using the pheatmap package (v1.0.10) in RStudio (1.1). Likelyhood Ratio Test was performed using the GTest function of the R package DescTools (v0.99). PCA analyses were performed using the PCA function of the FactoMineR package (v2.9). Relatedness of dendrogram based on gene expression and flow cytometry proportions were analyzed using the dendextend package (v1.19.1). Scripts used in the analysis and raw qPCR data are available at https://github.com/pgermon37/milk_RNA_profile.

## Results

### Identification of a gene panel able to discriminate macrophages, neutrophils and T lymphocytes

The aim of our study was to investigate if profiling of milk samples based on gene expression could provide relevant information regarding the inflammatory status of mammary gland quarters. For such an analysis to take into account the contribution of different leucocyte populations to the gene expression profile of raw milk samples, we first needed to select a panel of genes that showed different levels of expression depending on the cell type.

Therefore, we sorted macrophages neutrophils (live CD45+ G1+ cells), (live CD45+ G1-CD14+ cells), and T lymphocytes (live CD45+ CD3+ cells) by flow cytometry from four quarter milk samples having somatic cell counts ranging from 45,000 to 389,000 cells/ml (**Supplementary Table S1**).

The expression by these purified sorted cells of a panel of 47 genes was investigated by high-throughput RT-qPCR. These 47 genes were selected based on their expected role in the function of cell types studied, on the laboratory’s expertise in bovine immunology and on results of preliminary experiments on a larger set of 96 genes. Of the 47 genes selected, we identified 37 genes that were significantly differentially expressed between cell types (Kruskal-Wallis test - p < 0.05). Expression results for these 37 genes in the three selected cell types were then represented in a heatmap after hierarchical clustering (**Fig 1A**) and Principal Component Analysis (**Fig 1B**). Both analyses showed clear separation of samples based on cell types. The hierarchical clustering analysis allowed the identification of three clusters of genes with specificities for one of the cell types. The list of genes from these three clusters are presented in **Supplementary Table S3**.

**Fig 1:**
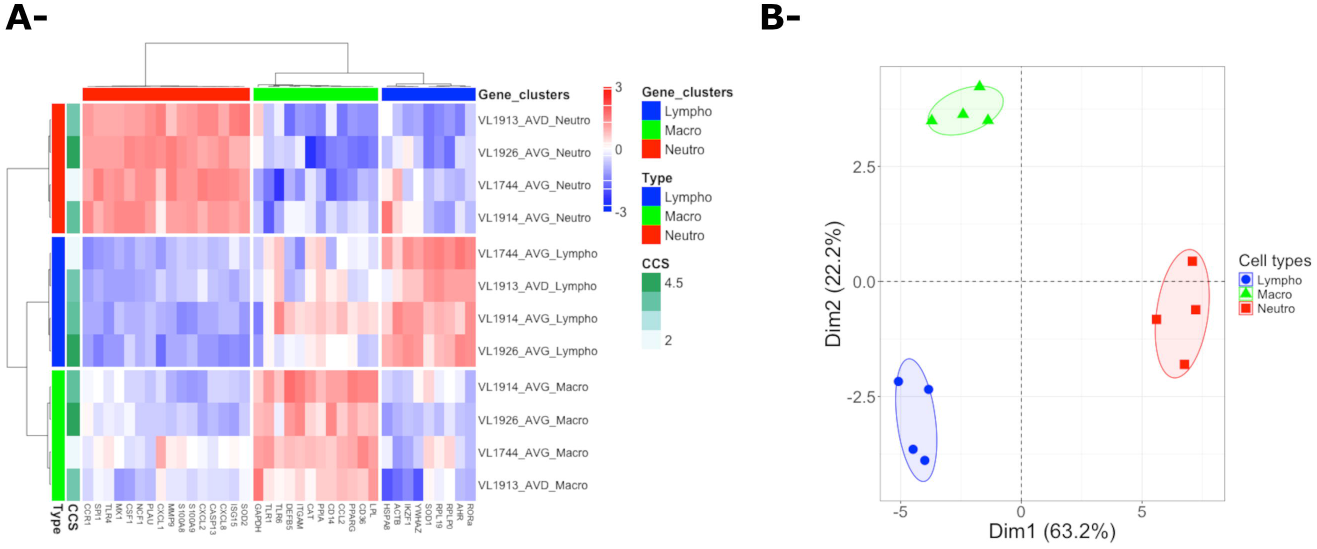
Heatmap and PCA of gene expression of sorted milk macrophages, neutrophils and T lymphocytes. Cells from four quarter milk samples from four different cows (one sample per cow) were collected by centrifugation, labelled with antibodies against CD3 (T lymphocytes), CD14 (macrophages) and G1 (neutrophils) and sorted on a MoFlo Astrios EQ cell sorter. RNA was extracted from purified cell populations and analyzed by high-throughput RT-qPCR. Gene expression values were calculated against a theoretical control sample. Genes significantly differentially expressed (p<0.05) among cell types were identified by performing a Kruskal-Wallis test and then analyzed by hierarchical clustering (A) and principal component analysis (B).

### Comparison of clustering based on gene expression and flow cytometry profiles

Based on these results, we investigated the expression profiles of these 37 genes in RNA extracted from raw milk samples. We analyzed quarter milk samples from 28 Holstein (HF) cows, which included the four quarter milk samples used for sorting the purified cell types, and from 10 Normande (NO) cows. One sample per cow was selected so that the panel of samples covered a range of SCC values from 17,000 to 3,603,000 cells/ml, with only 6 samples above the 400,000 cells/ml threshold. Cows were at different stages of lactation (days-in-milk ranging from 35 to 391 days). Detailed description of the different milk samples are given in **Supplementary Table S1**.

SCC distribution of selected milk samples depending on breed are depicted in **Supplementary Figure S2**: we could not detect any statistically significant difference in terms of SCC between samples from Holstein and Normande cows (Mann-Whitney p value = 0.29). The breed criteria was thus not explored further in downstream analyses.

For each sample, the gene expression profile after RNA extraction was determined by high-throughput RT-qPCR. Among the 37 genes selected above, expression values were obtained for all samples for 34 genes. Analysis by hierarchical clustering of expression values for these 34 genes, performed without any a priori, clearly identified four clusters of samples (**Fig 2A**). The PCA analysis showed a clear separation of these four clusters (**Fig 2B)** and a statistical analysis highlighted genes differentially expressed between clusters (**Fig 2C**). In particular, cluster 4 samples differ from clusters 2 and 3 mostly by differential expression of neutrophil-associated genes such as CXCL8, SOD2, TLR4, NCF1, CXCL2, CCR1 and SPI1. Cluster 1 and 2 samples differ from clusters 3 and 4 by a higher expression of lymphocyte-associated genes such as CD36, LPL, RORa, YWHAZ and PPIA.

**Fig 2.**
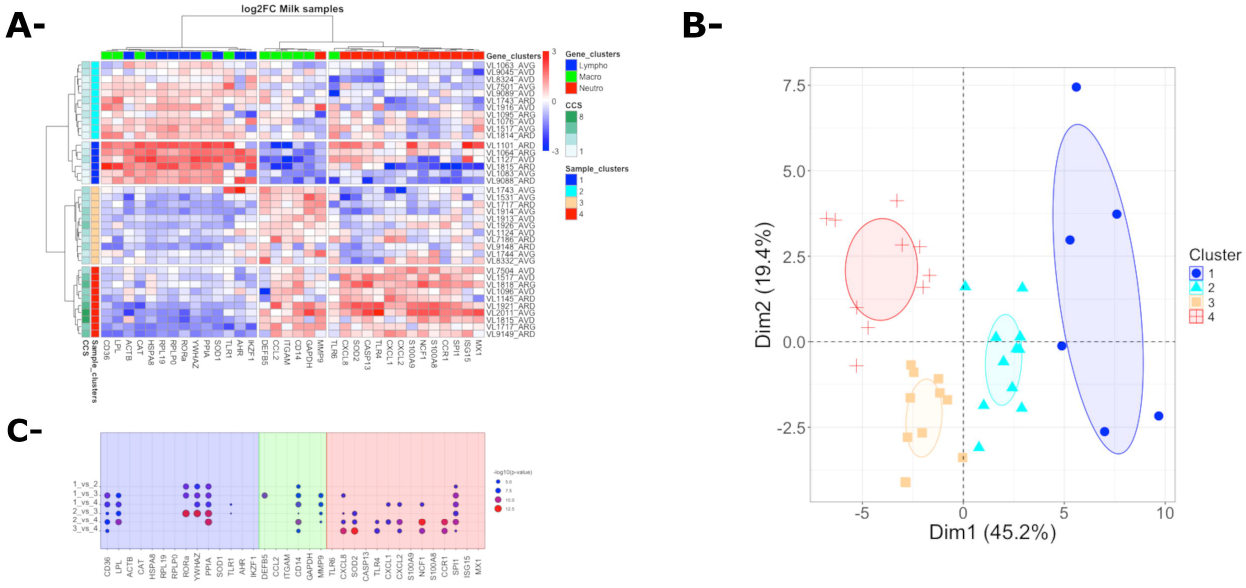
Heatmap and PCA of gene expression of raw milk samples. Milk samples were aseptically collected from 38 individual quarters. RNA was extracted directly from raw milk samples and analyzed by high-throughput RT-qPCR. Gene expression values were calculated against a theoretical control sample and analyzed by hierarchical clustering (A) and principal component analysis (B). Statistical significance of gene expression differences between clusters were first analyzed by Kruskal-Wallis tests followed by two-by-two comparison using Mann-Whitney tests (C). Only statistically significant differences are indicated (p<0.05). Color code for clusters is indicated on the figure.

Consistent with the higher expression of neutrophil-associated genes, cluster 4 samples are the ones with the highest SCC values while, even though they show different gene expression profiles, cluster 1 and 2 samples are both characterized by SCC values generally below 100,000 cells/ml (**Fig 3**). We could not detect any statistically significant difference in lactation stage between the different clusters (**Supplementary Figure S3** – Kruskal-Wallis p value = 0.35).

**Fig 3.**
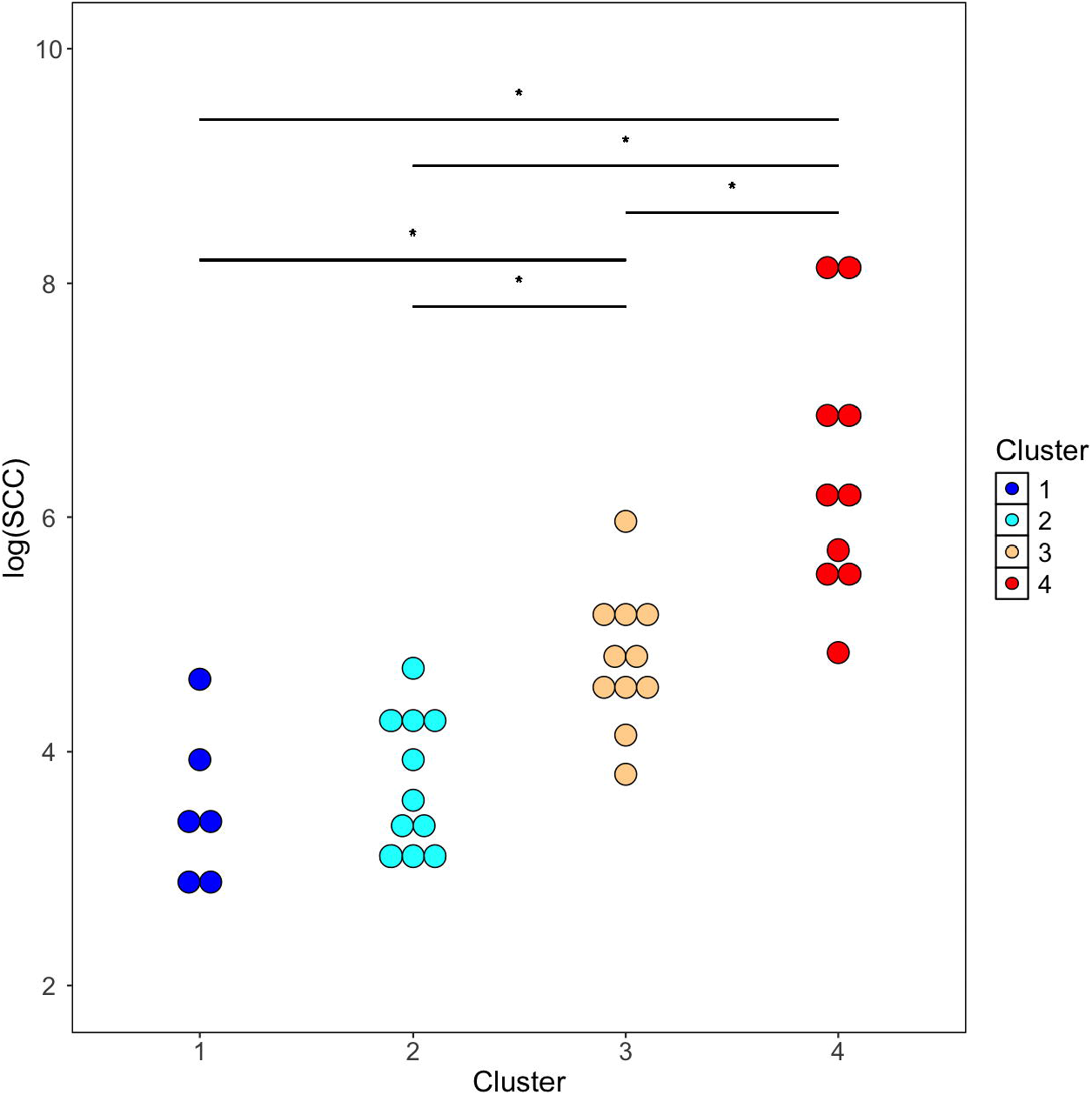
log(SCC) values of milk samples from the four different clusters. The log(SCC) of each of the milk samples analyzed is represented and samples are grouped and colored depending on the cluster to which they were allocated in the hierarchical clustering analysis. Color code is identical to that of Figure 2. * indicates statistical significance (p < 0,05). p-values were calculated using Mann and Whitney after global comparison using a Kruskal-Wallis test.

For each of these milk samples, the proportion of macrophages, neutrophils and T lymphocytes were obtained by flow cytometry (see **Supplementary Table S1**). Except for a few samples, analysis of these results by Principal Component Analysis showed a good overlap between the clustering based on gene expression profiles and that based on flow cytometry data (**Fig 4**).

**Fig 4.**
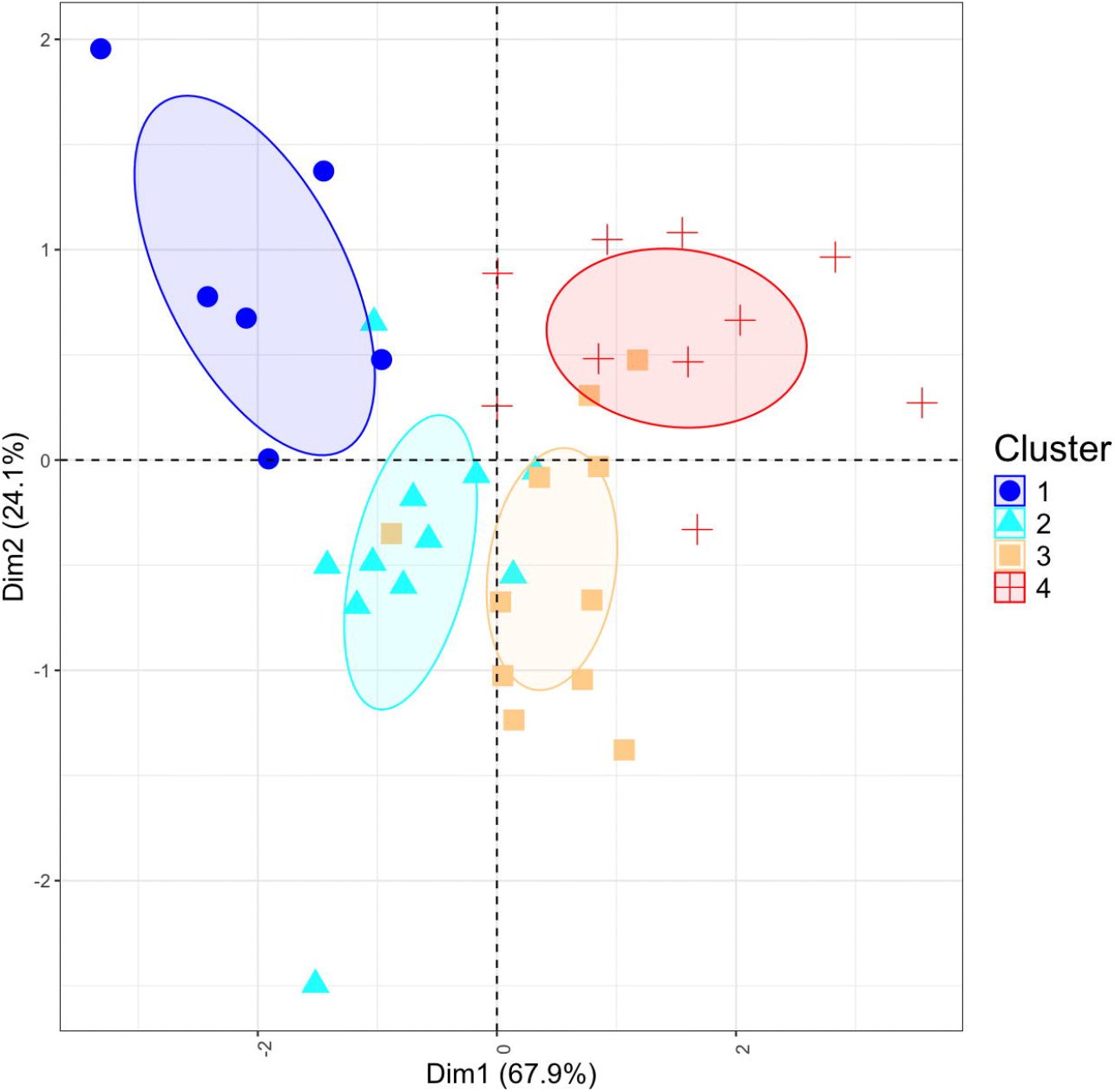
Principal component analysis of cellular composition of milk samples analyzed by flow cytometry. Counts of macrophages, neutrophils and T-lymphocytes among cells from collected milk samples was determined by flow cytometry. Principal Component Analysis of the cell composition was performed on the count values for each of the three cell types analyzed obtained by multiplying the SCC value by the percent of each cell type. Dots are colored depending on the cluster to which they were allocated based on the hierarchical clustering analysis of gene expression data. Color code is identical to that of Figure 2 and is indicated on the figure.

To quantitatively estimate the relatedness of analyses based on gene expression and flow cytometry, we analyzed the relatedness of dendrograms obtained with both methods calculating Baker’s index (data not shown). This analysis showed that the two trees share significant similarities: the null hypothesis that the two trees are not similar can be rejected with a risk below 0.01.

### Clustering based on gene expression profile differentiates samples with similar SCC

Classical bacteriological analysis was performed on all selected milk samples. Absence of bacteria was observed in 13 samples while, when bacteria were present, only Gram-positive pathogens were detected (16 samples). Bacteriological status could not be determined for 5 samples and 4 samples could not be examined due to technical reasons (**Supplementary Table S1**). When the clustering was analyzed relative to the presence of bacteria in the milk, the bacteriological status was significantly associated to the cluster type (Likelihood Ratio Test p-value = 0.015). In particular, the number of “No growth” sample is increased in cluster 1 samples while the number of samples showing growth of “Gram-positive” bacteria increases in cluster 4 samples (**Fig 5**).

**Fig 5.**
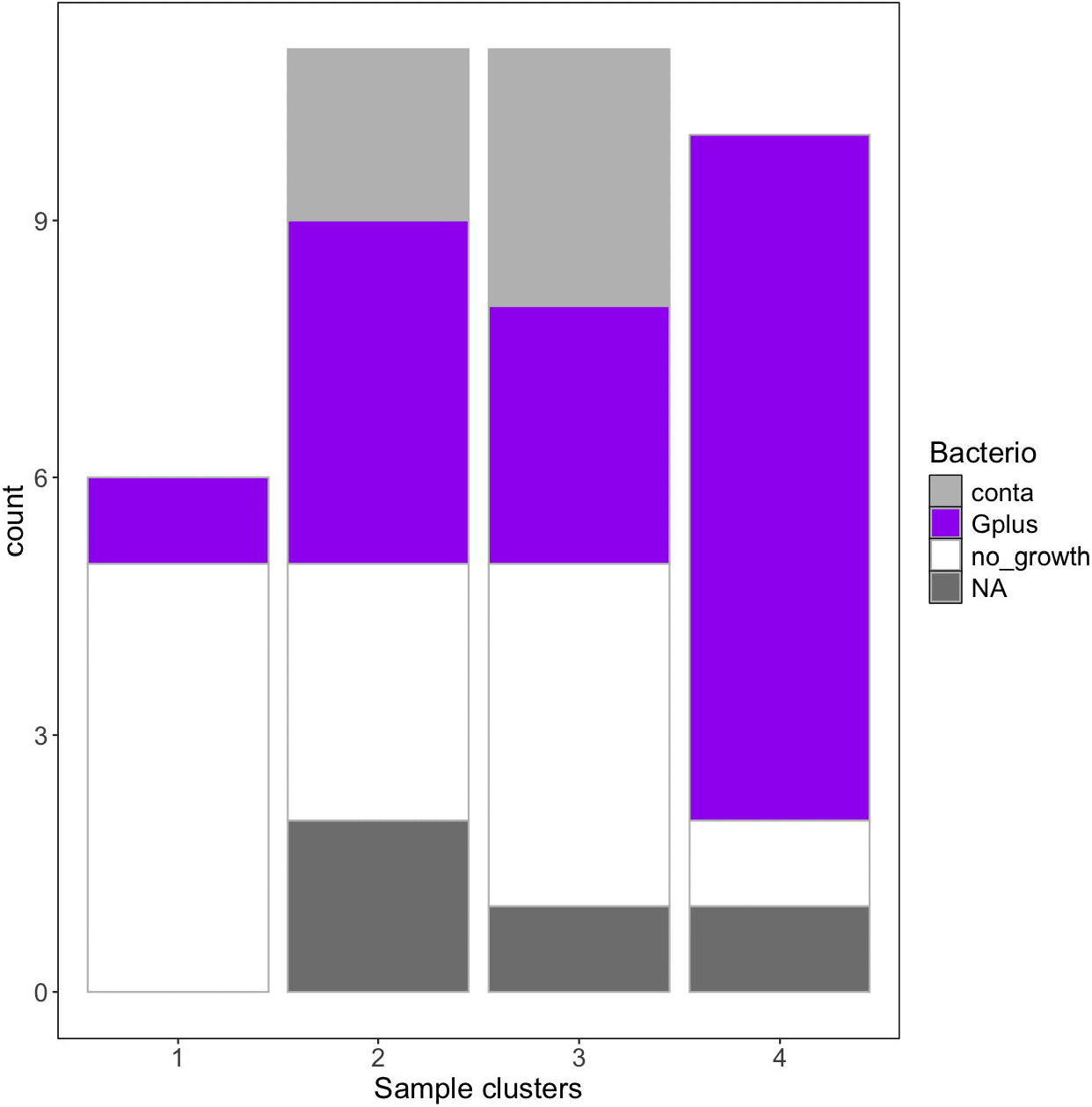
Barplot representing the number of samples in bacteriology positive or negative samples. Bacteriological status of each milk sample was analyzed by plating 50 microliter of each milk sample on sheep-blood agar plates. Samples with more that 3 types of colonies were considered as contaminated. Gram staining was performed on colonies growing on plates. NA indicates absence of bacteriological analysis.

## Discussion

The objective of the present study was to develop a new method for the functional characterization of the inflammatory status of mammary gland quarters. As an alternative to the characterization of cells present in milk, we investigated the gene expression profile of milk samples as a surrogate for the functionality of cells present in milk.

In order to determine if this strategy would allow us to characterize the inflammatory status of mammary glands, we first identified a subset of 37 genes that were differentially expressed between three cell types collected from milk, namely macrophages, T lymphocytes and neutrophils.

We then focussed our milk RNA expression analyses on low cell count milk samples, for which current strategies provide low positive predictive values of being infected: the median value for SCC of milk samples analyzed was 95 500 (Q1-Q3 : 38 250 – 218 000). In order to improve the diagnostic of inflammatory quarters, analyses were done on quarter milk samples. Indeed, analyses of composite samples show reduced predictive power compared to analyses based on quarter milk samples (35, 36). Importantly, it should be noted that none of the milk samples analyzed were from clinical cases, even the two samples with the high SCC values.

Based on the expression level of 34 genes from which expression could be measured in all samples, we identified four different clusters of milk samples, each with specific gene expression profiles. Cluster 4 samples are characterized by a higher expression of neutrophil-associated genes which is consistent with their high SCC values. Compared to samples from clusters 1 and 2, cluster 3 and 4 samples are typified by increased expression of macrophage related genes and lower expression of genes typifying T lymphocytes. Most interestingly, as can be seen in **Figure 2**, our method was able to differentiate cluster 1 and 2 samples despite their similarly low SCC values in the range of 50,000 to 200,000. These samples are both characterized by a higher level of expression of lymphocyte-associated genes compared to clusters 3 and 4. Yet, cluster 1 samples show a higher expression of some lymphocyte-associated genes, a lower expression of CD14, a macrophage-associated gene, and a higher expression of SPI1. These differences in gene expression suggest different immune status between these cluster 1 and 2 samples.

Due to the lack of functionnal tests and the somehow limited number of genes studied, we can only speculate on the link between these expression profiles and the inflammatory status of mammary gland quarters. For instance, because of a reduced expression of neutrophil-associated genes and an increased expression on macrophage-related genes, we could hypothesize that cluster 3 samples originate from mammary gland quarters in the process of resolution with increased proportions of dead neutrophils and higher macrophage counts. Alternatively, they could also be representatives of chronically infected cows or be specific to a particular pathogen. We could also hypothesize that cluster 1 samples correspond to healthy samples with limited inflammation. Compared to cluster 2 samples, cluster 1 samples have higher expression levels of macrophage-associated genes: considering these samples as healthy is consistent with the observation of higher proportions of macrophages in healthy mammary glands (9, 25, 26).

The method described in this pilot study has several advantages over current methods to qualify the inflammatory status of mammary gland quarters. Compared to clustering strategies proposed in other studies, such as grouping samples based on SCC values and bacteriological status or on SCC and DSCC values (11, 37), the clusters we have identified were obtained without any a priori, are only based on intrinsic properties of the milk samples linked to macrophages, neutrophils and T lymphocytes and are not conditioned by user-defined thresholds. It should be noted that even in low SCC milk samples a clear gene expression pattern could be observed. Therefore, contrary to DSCC which requires SCC values above 50,000 cells/ml, a low SCC is not a limit to the use of our method (25).

In addition, the protocol presented here does not rely on preliminary centrifugation of milk samples which, as can be the case for flow cytometry methods, might fail to recover all cells; in particular macrophages that can remain in the cream layer after centrifugation (9).

We therefore believe that it could bring an added value to the definition of inflammatory status of mammary gland quarters. More specifically, this method could be used to investigate in more details the response of mammary gland quarters to infection. For instance, neutrophil subtypes in different inflammatory conditions could be investigated which would extend our observations of increased proportions of a neutrophil subset expressing MHCII surface molecules during subclinical mastitis (38). Moreover, in vaccine based studies were a local immune response is expected, the gene panel could be adapted to monitor more precisely the response of T-lymphocytes such as the Th17 response that has been shown to be beneficial to the clearance of bacteria (39, 40).

For these more detailed analyses to be possible, new developments will be required to increase the panel of genes analyzed in order to better distinguish additional cell types or subtypes. Single-cell RNAseq analyses have recently been performed on milk cells (33). These data confirmed some of the associations described herein: for example, CXCL8 and TLR4 were also identified as specific for granulocytes/neutrophils. CD14 was identified as associated to macrophages, as expected for a macrophage specific gene (41). More detailed analyses of these data and of the cell clusters observed could provide usefull information regarding additionnal genes specific for different cell types or even subtypes.

The method described herein nevertheless has certain limitations. The main limit is that it is not easily accessible to on-farm diagnostic of mammary gland inflammation and requires specific equipments. Its interest is limited for the moment to laboratory studies trying to better qualify the inflammatory status of the udder. We also failed to clearly quantify epithelial cells by flow cytometry. Although these cells are likely to represent only a low proportion of milk cells, their proportions could potentially modify the clustering observed by flow cytometry. Nevertheless, it is likely that the gene expression profiling of milk samples is actually susceptible to the presence of epithelial cells in milk (42). It is also possible, since fat globules have been shown to contain RNA molecules originating from the milk producing epithelial cell, that the gene expression profile also reflects the functionality of mammary epithelial cells (43). As a consequence, RNA quantification from raw milk might also reflect both the presence and/or functionality of epithelial cells.

The present report should be considered as a pilot study that was limited to only a small number of samples. In order to validate the usefulness of our method, more broader studies would be required. A more detailed investigation of the relationship between gene expression clusters and the bacteriological status of mammary gland quarters would definitely support the predictive value of these analyses. So far, only an tendency for an increased proportion of samples with Gram+ bacteria in cluster 4 samples was observed, consistent with the fact that these samples are the ones with the highest SCC. It also remains to be determined whether the gene expression profile varies depending on parity, days-in-milk as these parameters are clearly associated with variations in SCC and DSCC (44). Complementary analyses should also take into account the variability of milk cellular composition at different stages during milking (45). The current profiles identified were based on foremilk samples which are likely to be enriched in SCC in samples with more than 100,000 cells/ml (46).

To conclude, we set up a new way of monitoring the inflammatory status of mammary gland quarters during lactation based on the profiling of gene expression from raw milk with minimal processing before RNA extraction. We were able to distinguish four types of milk samples that corresponded to different inflammatory status. Potential refinements include the addition of new genes to better distinguish quarters with low SCC values, improved qualification of the inflammatory status of mammary glands along with more in-depth characterization of bacteria present in milk samples.

Although not applicable as such in commercial farms, this method paves the way to the development of new medium- or high-throughput phenotyping strategies. Such methods will provide useful information for the implementation of selective dry cow therapy or for the routine analysis of health status of mammary glands.

## Acknowledgements

The authors would like to thank the animal staff of the “Unité Expérimentale du Pin-au-Haras” (INRAE, Gouffern-en-Auge, France) and of the “Unité Expérimentale de Physiologie animale de l’Orfrasière” (INRAE, Nouzilly, France) for their help in collect ing milk samples. We are grateful to Aude Remot for critical reading of the manuscript.

## Data, scripts, code, and supplementary information availability

Data are available online: https://doi.org/10.5281/zenodo.20056104 Supplementary information is available online: https://doi.org/10.5281/zenodo.20281966

