## Supplementary figures and images for "Development of a new tool based on gene expression analyses of bovine raw milk to monitor the inflammatory status of the udder"

### S1_Fig.tif

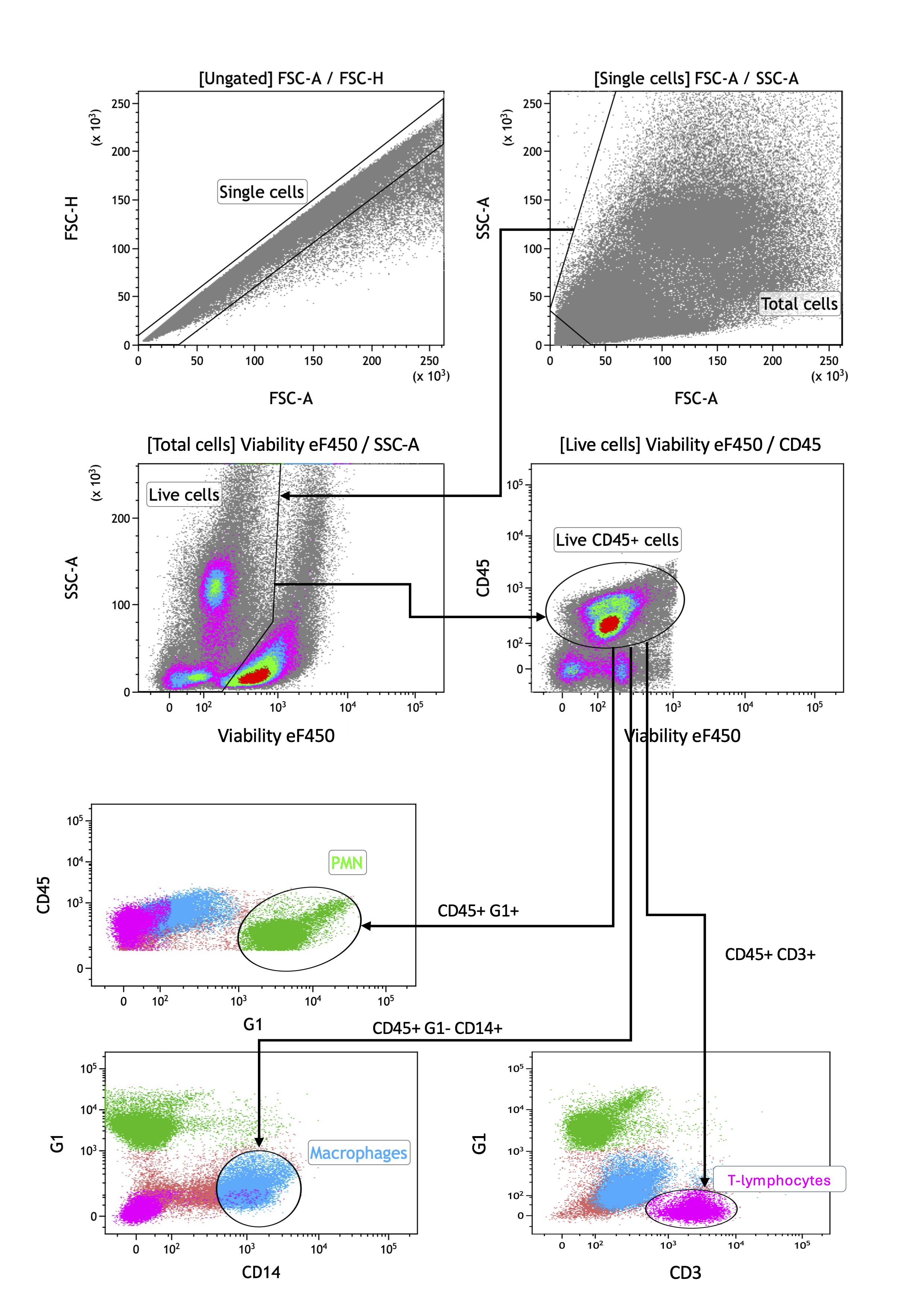

### S2_Fig.tif

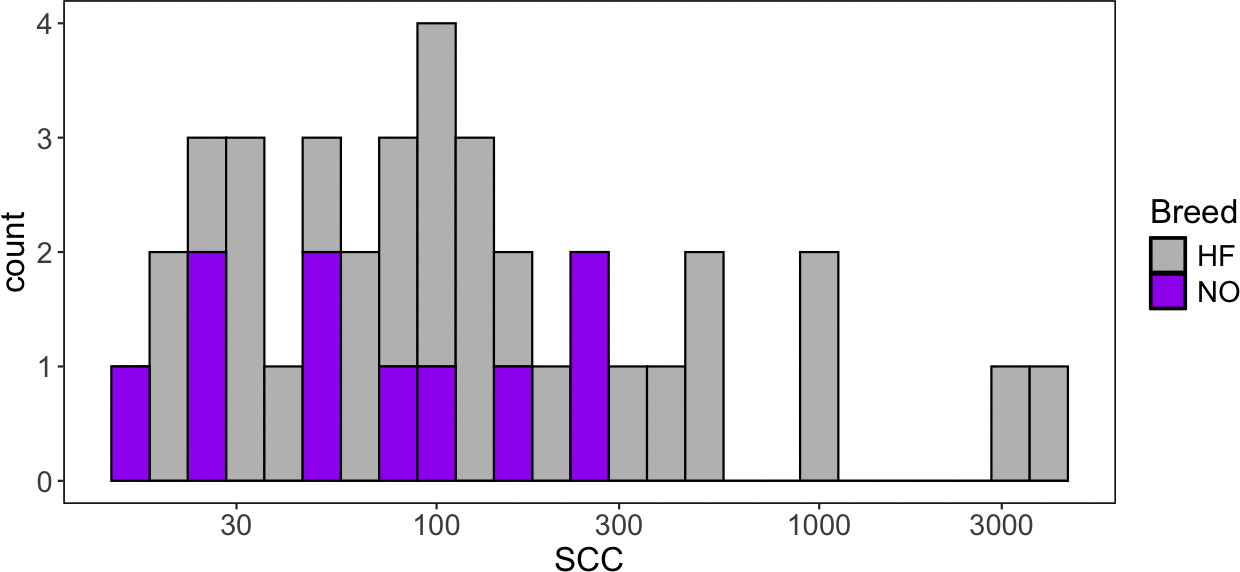

### S3_Fig.tif

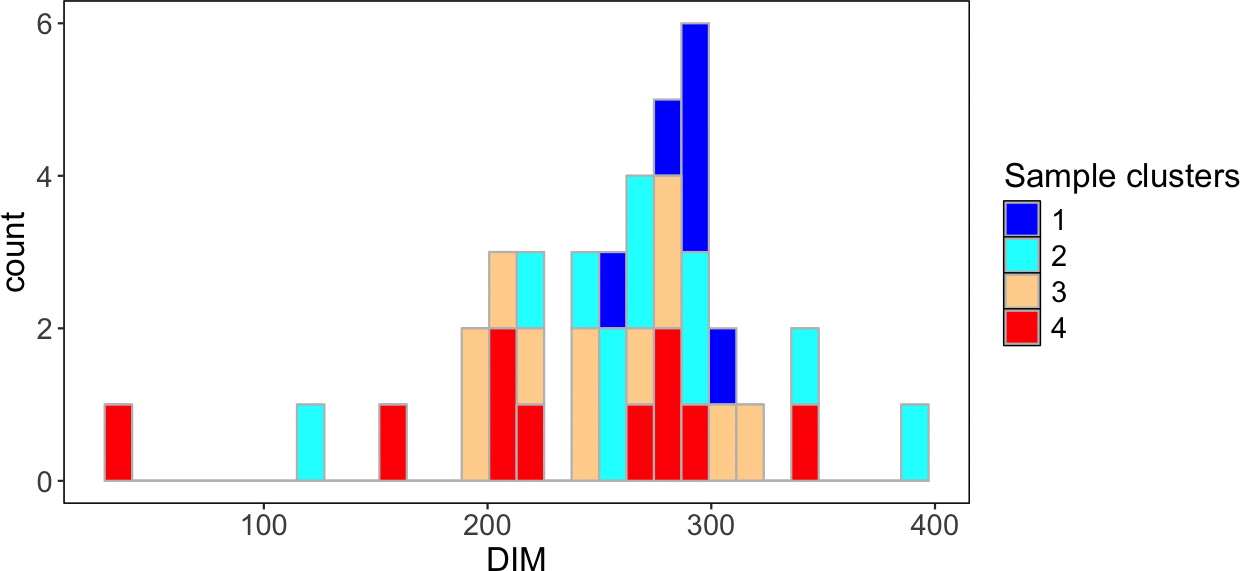
